# An eQTL landscape of kidney tissue in human nephrotic syndrome

**DOI:** 10.1101/281162

**Authors:** Christopher E. Gillies, Rosemary Putler, Rajasree Menon, Edgar Otto, Kalyn Yasutake, Viji Nair, Paul Hoover, David Lieb, Shuqiang Li, Sean Eddy, Damian Fermin, Nephrotic Syndrome Study Network (NEPTUNE), Nir Hacohen, Krzysztof Kiryluk, William Wen, Matthias Kretzler, Matthew G. Sampson

## Abstract

Expression quantitative trait loci (eQTL) studies illuminate the genetics of gene expression and, in disease research, can be particularly illuminating when using the tissues directly impacted by the condition. In nephrology, there is a paucity of eQTLs studies of human kidney. Here, we used whole genome sequencing (WGS) and microdissected glomerular (GLOM) & tubulointerstitial (TI) transcriptomes from 187 patients with nephrotic syndrome (NS) to describe the eQTL landscape in these functionally distinct kidney structures.

Using MatrixEQTL, we performed *cis*-eQTL analysis on GLOM (n=136) and TI (n=166). We used the Bayesian “Deterministic Approximation of Posteriors” (DAP) to fine-map these signals, eQtlBma to discover GLOM-or TI-specific eQTLs, and single cell RNA-Seq data of control kidney tissue to identify cell-type specificity of significant eQTLs. We integrated eQTL data with an IgA Nephropathy (IGAN) GWAS to perform a transcriptome-wide association study (TWAS).

We discovered 894 GLOM eQTLs and 1767 TI eQTLs at FDR <0.05. 14% and 19% of GLOM & TI eQTLs, respectively, had > 1 independent signal associated with its expression. 12% and 26% of eQTLs were GLOM-specific and TI-specific, respectively. GLOM eQTLs were most significantly enriched in podocyte transcripts and TI eQTLs in proximal tubules. The IGAN TWAS identified significant GLOM & TI genes, primarily at the HLA region.

In this study of NS patients, we discovered GLOM & TI eQTLs, identified those that were tissue-specific, deconvoluted them into cell-specific signals, and used them to characterize known GWAS alleles. These data are publicly available for browsing and download at http://nephqtl.org.

## Introduction

Nephrotic syndrome (NS) is a rare disease of glomerular filtration barrier failure^1;2^ causing massive urinary excretion of protein, that can progress to chronic kidney disease (CKD) and end-stage renal disease (ESRD)^3;4^. As NS is a heterogeneous disease, we use the histologic descriptions of glomeruli on kidney biopsy to diagnose patients with “minimal change disease (MCD)” and “focal segmental glomerulosclerosis (FSGS)”. Or we use a patient’s response to these treatments to give them a *post hoc* diagnosis of steroid sensitive NS (SSNS) or steroid resistant NS (SRNS).

Understanding how human genetic variation contributes to the development and progression of NS has been a fruitful strategy in gaining a more precise understanding of the molecular underpinnings of NS^5^. More than 50 genes have been discovered that harbor rare variants sufficient to cause SRNS (“Mendelian” NS)^6^. Through genome-wide association studies (GWAS) and exome-chip studies^7-10^, common genetic variants have been discovered that contribute to the pathogenesis of FSGS, pediatric SSNS, and membranous nephropathy (MN). Rare variant association studies in FSGS have implicated a small set of genes harboring an increased burden of rare, deleterious variants^8^. We are challenged to discover addition forms of NS-associated genetic variation to gain biological and clinical insights.

Expression quantitative trait loci (eQTL) studies, which use mRNA expression as a proximal (and continuous) cellular endophenotype, have increased power for discovery of statistically significant genetic effects as compared to GWAS and provides inherent biological meaning in the associations between a regulatory variant and its associated gene^11-14^. The GTEx project has generated eQTL data which is publicly available and has been used extensively to help interpret GWAS signals emerging from other complex traits^15^. Meaningful eQTL discoveries using the affected tissues in other human diseases suggest their potential for NS genomic discovery as well. This is appealing given that we often obtain kidney tissue via biopsy from affected patients.

With regards to kidney eQTLs, the final release of GTEx will only have 73 kidney cortex samples (*Wen, personal communication)*. There is also an absence of any other major public kidney eQTL datasets. This represents a major barrier for genomic discovery in nephrology.

The most comprehensive kidney eQTL study thus far discovered kidney eQTLs using unaffected portions of 96 nephrectomy samples from The Cancer Genome Atlas^16^. The investigators integrated these eQTL with risk loci for chronic kidney disease (CKD) to establish links between these risk alleles and molecular mechansims^16^. A limitation of this study was that bulk renal cortex was used for association, which is known to be 80% proximal tubule cells. The preponderance of this cell type may obscure eQTL signals emerging from structurally and cellularly heterogeneous kidney. Another difference is that this study exclusively used healthy tissue, which prevents an opportunity to potentially discover disease-context specific eQTL effects.

Microdissecting bulk renal cortex tissue into its two main functional structures, the glomerulus (GLOM) and tubulointerstitium (TI), allows increased specificity for kidney transcriptomics studies. For instance, targeted GLOM and TI eQTL studies have led to discoveries of the transcriptomic impact of diabetic kidney disease GWAS alleles^17-19^ in patients with diabetic nephropathy. In NS, a GLOM eQTL study of apolipoprotein L1 (*APOL1*) high-risk genotype^20^ led to a number of insights which have been subsequently followed up genetically and experimentally^21^.

The success of previous tissue-specific eQTL studies in affected populations motivate us to use this strategy for NS genomic discovery. Here, we report the results of a genome-wide GLOM and TI cis-eQTL study from 187 biopsied NS patients enrolled in the Nephrotic Syndrome Study Network (NEPTUNE)^22^. In addition to this report, we have also created “nephQTL,” a publicly available eQTL browser to share these data with the wider community (http://nephqtl.org).

## Materials and Methods

### Data Sources and Patient Inclusion

The Nephrotic Syndrome Study Network (NEPTUNE) is a prospective, longitudinal cohort recruiting participants with substantial proteinuria at the time of first kidney biopsy for clinical suspicion of minimal change disease (MCD), focal segmental glomerulosclerosis (FSGS), or membranous nephropathy (MN)^22^. Phenotypic data, urine, and blood samples are collected at baseline and over time. DNA was collected for a variety of genotyping studies and a research renal biopsy core was collected for transcriptomic analysis^20;23-25^.

### Whole-genome sequencing

On 322 NEPTUNE subjects, we used an Illumina Hi-Seq to perform low-depth WGS, which takes advantage of shared haplotypes to make accurate genotype calls^26^. We used GotCloud^27^ as our standard pipeline for alignment and variant calling (including insertion-deletions [indels]). A subset of patients also underwent Illumina Exome Chip SNP genotyping. Using Exome chip genotypes as our gold standard, we calculated their concordance with non-monomorphic sites on WGS at 77,769 shared sites. 97.5% of sites had > 97% concordance. Site-level genotype data from WGS and the Exome Chip, summarized across this cohort, are available at http://nephvs.org.

### Gene Expression

Gene expression data from microdissected GLOM and TI was generated using Affymetrix 2.1 ST chips^28^ and quantified with a Custom CDF file from BrainArray for EntrezG, version 19^29^. Expression was normalized across genes using robust multi-array average (RMA)^30^. PEER factors were computed as previously described^31^.

### Patient Inclusion

There were 197 NEPTUNE participants eligible for this study because they had WGS data, transcriptomic data for at least GLOM or TI, and clinical and demographic information. There were 136 and 166 available for GLOM and TI eQTL discovery, respectively. Of these, 115 patients were part of both the GLOM and TI eQTL analyses (**Figure 1**).

**Figure 1:**
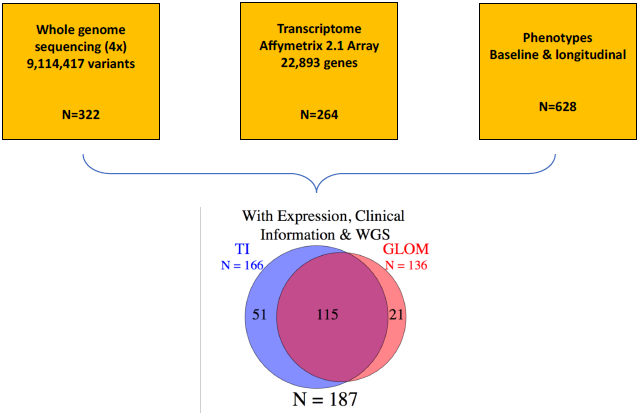
Datasets used in the present study and the number of eligible participants as a function of their available biopsy-based transcriptomic datasets.

### eQTL Analysis

We used MatrixEQTL^32^ for the initial step in the eQTL study. We focused solely on *cis* eQTLs only because we lacked substantial power for *trans* studies. Eligible variants were those with an AF>0.03 in our cohort and within 500 kb of qualifying genes’ transcription start or end sites (and within the gene itself). We adjusted for age, sex, the first four principal components of genetic ancestry, and PEER factors. Principal components of genetic ancestry were calculated using EPACTS^33^ on LD-pruned whole genome sequence data across all 187 individuals for whom we had expression information **(Supplementary Figure 1)**. We adjusted for the first four PCs, which accounted for a majority of the population variation. PEER factors were created utilizing the PEER framework as previously described^31;34^, adjusting for patient age, sex, and microarray batch. We adjusted for 31 PEER factors in GLOM and 25 in TI, the number that maximized significant eQTLs at an arbitrary p-value threshold and portion of the genome. To control the FDR at gene-level from the MatrixEQTL output, we used TORUS^35^, which provides computationally efficient estimation of Bayesian priors.

### Fine Mapping with DAP

We utilized the estimated priors from TORUS, accounting for variant distances to TSS of the target genes, to perform variant fine-mapping of eQTLs for each gene, using the Deterministic Approximation of Posteriors (DAP) algorithm^36^. DAP identifies independent signals contributing significantly to changes in expression, and assigns a posterior inclusion probability (PIP) to them. DAP incorporates functional genomic annotations, and accounts for patterns of linkage disequilibrium (LD) among SNVs.

Independent association signals were either single SNVs or indels or groups of them in LD (r^2^ > 0.25). The significance of each signal is characterized by local FDR. Those with local FDR < 0.05 were considered significant association signals in which we could predict the driving variants. Due to high LD and similar effect, the exact variant that drives the association signal is not always easily determined. DAP provides within-signal posterior inclusion probabilities (PIP) for each variant to help identify the most likely driver. The member SNVs for each signal group naturally form a 95% Bayesian credible set.

### Tissue specific mapping with eQtlBma

To assess the extent of tissue specificity, we used eQtlBma to estimate the proportion of eQTLs shared across GLOM and TI^37^, using summary statistics generated by MatrixEQTL. This framework also has the benefit of maximizing power over tissue-by-tissue analysis by jointly analyzing the tissues and allowing for differences in eQTLs by tissue, and improving power for detection of shared eQTLs.

### Geneset enrichment analysis

To better understand the biologic context of the genesets emerging from our DAP and eQTLBMA analyses, we used the Genomatix Software Suite (Genomatix; Munich, Germany) and Ingenuity Pathway Analysis (Qiagen, Inc)^38^ for functional annotation. In the IPA analysis, a complex network was constructed among the pathways that shared three or more genes to highlight “clusters” and pathways that crosstalk between the clusters.

### Generation of single cell RNA-Seq data

We combined and analyzed three scRNA-Seq datasets generated from healthy portions of tumor-nephrectomy samples specifically harvested for single cell analysis using 10X Genomics methodology. Single cell dissociations were done as reported39. Individual cells were labeled with barcodes, and transcripts within each cell were tagged with distinct UMIs (Unique Molecular Identifiers) in order to determine absolute transcript abundance. The library quality was assessed with “High sensitivity cDNA arrays” on an Agilent BioAnalyzer (ThermoFischer) platform. Sequencing was done on Illumina HiSeq2500 with 2 × 75 paired-end kits using the following read lengths: 26 bp Read1 and 110 bp Read2.

The sequencing data was first analyzed using the Cell Ranger software (10x Genomics Inc., Pleasanton, CA, USA) in order to extract the gene expression data matrix. Each element of the matrix is the number of UMIs associated with a gene and a barcoded cell. Next, we filtered out cells with less than 500 genes per cell and with more than 25% mitochondrial read content. Further downstream analysis steps used the Seurat R package include normalization, identification of highly variable genes across the single cells, scaling based on number of UMI and batch effect, dimensionality reduction (PCA, and t-SNE), standard unsupervised clustering, and the discovery of differentially expressed cell-type specific markers. Differential gene expression to identify cell-type specific genes was performed using the non-parametric Wilcoxon rank sum test.

### Cell-type deconvolution using kidney single cell RNA-Seq

We wanted to discover if our significant GLOM and TI eQTLs were enriched in specific kidney cell types. We began by analyzing our adult kidney scRNA-Seq data to identify genes whose expression were enriched in a particular cell-type. We then computed the enrichment of GLOM and TI eQTLs in these cell-type enriched genesets using a Fisher’s Exact test.

### Transcriptome wide association study with IgA Nephropathy GWAS

Previous studies have shown that risk alleles from genome-wide association studies (GWAS) are enriched for eQTLs^40^, and others have used eQTL data to pinpoint genes whose expression was regulated by GWAS alleles^16^. Here, we integrated our GLOM and TI eQTL data with summary statistics from the largest published GWAS of glomerular disease, in IgA Nephropathy^41^ (IgAN), and performed a transcriptome wide association study (TWAS) using an approach adapted from the PrediXcan and MetaXcan methods^42;43^.

Using the same genotype and expression data from the eQTL analysis, we first adjusted each gene’s expression by age, sex, four genetic PCs, and 31 PEER factors in TI and 25 in GLOM. Using the residuals for each gene we used the R package “glmnet” to fit a regression equation penalized using an elastic net with α=0.5, which is a mixture of an L1 and L2 penalty. For each gene we allowed non-ambiguous biallelic variants within 500kb of the start and end positions of each gene that were also present in the IgA GWAS and eQTL dataset (AF > 0.03 in the eQTL dataset). To select an appropriate hyperparameter, we used 30-fold cross-validation and selected the parameter that maximized the prediction R^2^ of the validation set. We selected genes with a cross-validated prediction R^2^ > 0.01.

Using a reference panel comprising East Asian and European samples from the 1000 Genomes data (n=1007 samples), for each gene *g*, we computed the variance of SNV l 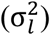 and variance of predicted expression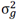. The variance 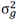 was defined in the MetaXcan paper as:

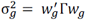

, where 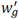 (is a vector of weights for gene *g*, and Γ is a covariance matrix of the SNPs included for gene g computed across the samples selected above.

Finally, we computed the *Z*_*g*_ statistic for gene g as:

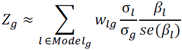

where *Model*_*g*_is the set of SNPs for gene *g, w*_*lg*_ is the weight learned for gene *g* and SNV *l* from the elastic net, β _*l*_ is the effect size from the GWAS result for SNV *l*, and 45(β_*l*_) is the standard error for the effect siz e. We performed gene-level association using only the genes for which we could predict expression with a cross-validated r^2^ > 0.01.

## Results

Characteristics of the included patients are presented in **Table 1**. Approximately 30% of patients had childhood onset of disease and the median duration of disease prior to biopsy was 4 months, suggesting that we were not studying the transcriptome of patients with longstanding proteinuria. The histologic diagnosed were almost equally divided between minimal change disease (MCD), focal segmental glomerulosclerosis (FSGS), membranous nephropathy (MN), and other glomerular conditions (most often IgA nephropathy).

**Table 1:** Patient characteristics for NEPTUNE participants included in eQTL analyses.

| **Characteristic** | **Either Expression (n=187)** | **GLOM Expression (n=136)** | **TI Expression (n=166)** |
| --- | --- | --- | --- |
| *Age (y)* | 36 (17-56) | 34.5 (17-55) | 36 (17-56) |
| *Pediatric* | 51 (27%) | 39 (29%) | 46 (28%) |
| *Age of Onset (y)* | 36 (17-56) | 34.5 (17-55) | 36 (17-56) |
| *Duration of disease (mos)* | 4 (1-18) | 4 (1-17.8) | 4 (1-18) |
| *Male* | 131 (70%) | 96 (71%) | 115 (85%) |
| *Histopath* |  |  |  |
| *FSGS* | 55 (29%) | 35 (26%) | 50 (30%) |
| *MCD* | 37 (20%) | 28 (21%) | 35 (21%) |
| *MN* | 41 (22%) | 36 (26%) | 35 (21%) |
| *Other* | 54 (29%) | 37 (27%) | 46 (28%) |
| *Clinical characteristics at baseline* |  |  |  |
| *RAAS blockade* | 130 (70%) | 94 (69%) | 117 (70%) |
| *eGFR* | 82.5 (56.4-105.6) | 83.8 (65.7-107.6) | 82.4 (54.4-105.5) |
| *UPCR* | 2.2 (0.8-4.2) | 2 (0.6-3.9) | 2.1 (0.8-3.9) |
| *Complete Remission* | 108 (58%) | 83 (61%) | 96 (58%) |

### Matrix eQTL

Using Matrix eQTL, we analyzed 76,979,158 *cis-*pairs across 9,114,417 SNV and expression from 22,893 genes. Based on the minimum FDR values across all eQTLs, there were 1055 genes with significant eQTLs in GLOM and 3217 in TI, at FDR < 0.05.

### DAP & eQtlBma

The output from MatrixEQTL plus the local FDR from TORUS were “fine-mapped” using DAP. Rather than solely identifying that an eQTL exists, DAP identifies eQTLs in which the specific variants or “clusters” of variants predicted to be driving the association can be confidently identified. In addition, DAP can discover eQTLs in which multiple independent SNVs or clusters of SNVs are associated with the gene’s expression.

Using DAP, we discovered 894 GLOM eQTLs and 1767 TI eQTLs at < 5% FDR level (**Supplementary Table 1**). The majority of eQTLs had one independent signal responsible for the association. Multiple independent signals associated with expression were found in 112 GLOM eQTL (14%) & in 337 in TI (19%). (**Supplementary Figure 2**).

**Figure 2:**
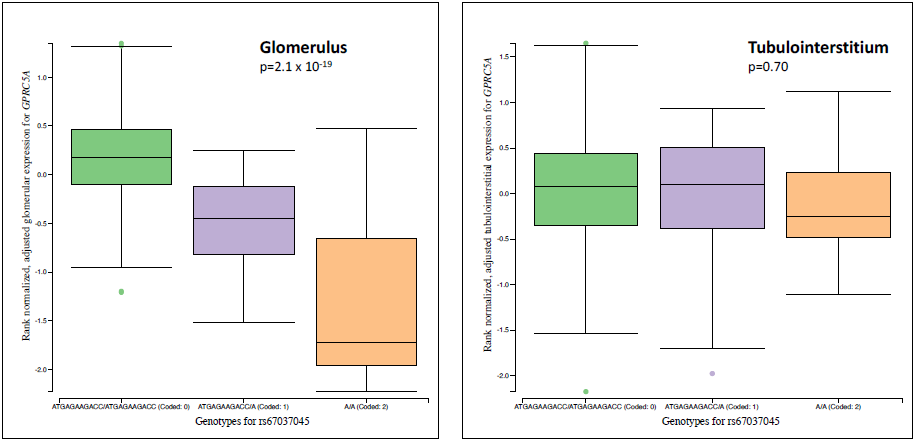
Matrix eQTL output for *GPCR5A*, among top-ranked glomerular specific eQTL as computed by eQtlBma.

To identify eQTLs that were specific to, or shared between, the GLOM & TI, we used eQtlBma, with the MatrixEQTL data as input. We estimated 12.2% and 26.3% of eQTLs are GLOM-specific and TI-specific, respectively (**Figures 2 & 3; Supplementary Table 2**). Note that this tissue specificity estimate of eQTLs is not a simple tally of individual eQTL signals. It is obtained by pooling all genes across the two tissues simultaneously, which takes the power difference in each tissue into account.

**Table 2:**
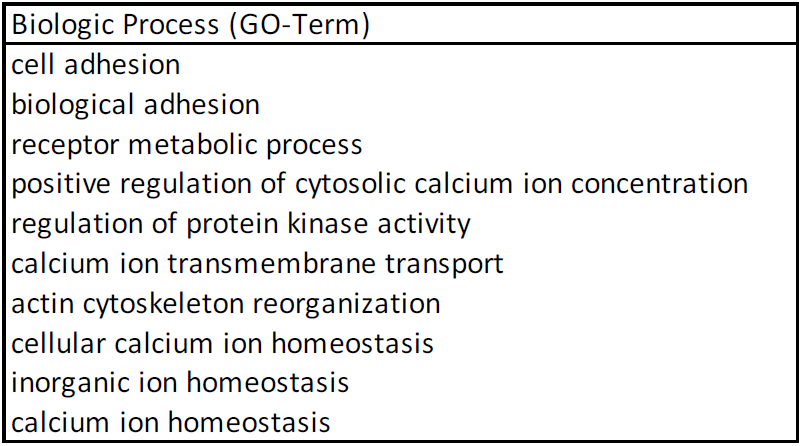
Top GO biologic processes for GLOM-specific eQTLs as defined by eQtlBma

| Biologic Process (GO-Term) |
| --- |
| cell adhesion |
| biological adhesion |
| receptor metabolic process |
| positive regulation of cytosolic calcium ion concentration |
| regulation of protein kinase activity |
| calcium ion transmembrane transport |
| actin cytoskeleton reorganization |
| cellular calcium ion homeostasis |
| inorganic ion homeostasis |
| calcium ion homeostasis |

We performed geneset enrichment analysis of the DAP-derived significant GLOM and TI eQTLs using Genomatix and IPA (Qiagen, Inc) software. Analyzing the IPA networks showed a number of immunity pathways in GLOM (**Figure 4a**). The TI pathways also were enriched for immunity; however metabolic and oxidative pathways were equally as enriched (**Figure 4b**).

A striking aspect of the pathway enrichment analyses was the shift from inflammatory/immune related pathways and processes in the DAP GLOM gene set to those more specific to podocyte biology with the eQTLBMA GLOM gene set. For example, 8 of the top 10 GO Biologic Processes for the DAP GLOM gene set were related to antigen presentation or interferon gamma signaling. By contrast, the top 10 GO Biologic Processes for GLOM eQTLs via eQTLBMA included actin cytoskeleton rearrangement, calcium signaling, and cell and biologic adhesion (**Table 2**). A StringDB network created from the eQtlBma GLOM-specific gene showed a network significantly enriched for interactions with the two most interconnected genes being integrin alpha V (*ITGAV)* and transforming growth factor beta 1 *(TGFB1*) (**Supplementary Figure 3**).

**Figure 3:**
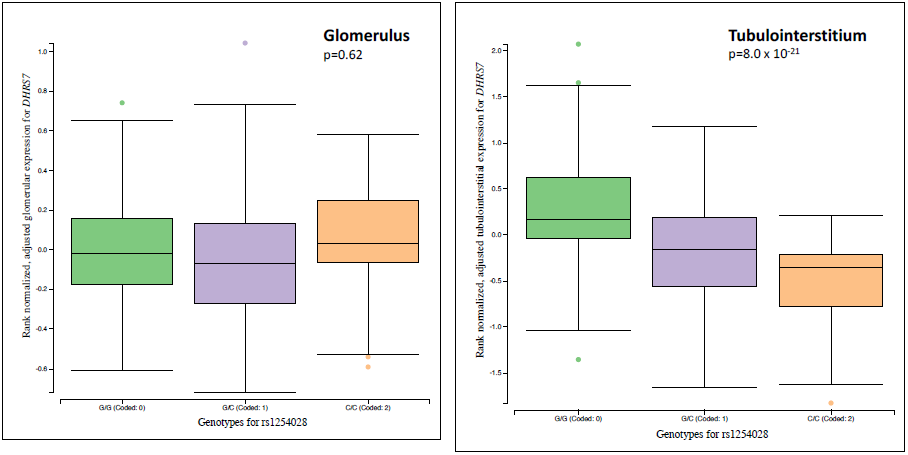
Matrix eQTL output for *DHRS7*, among top-ranked tubulointerstitial-specific eQTL as computed by eQtlBma.

**Figure 4:**
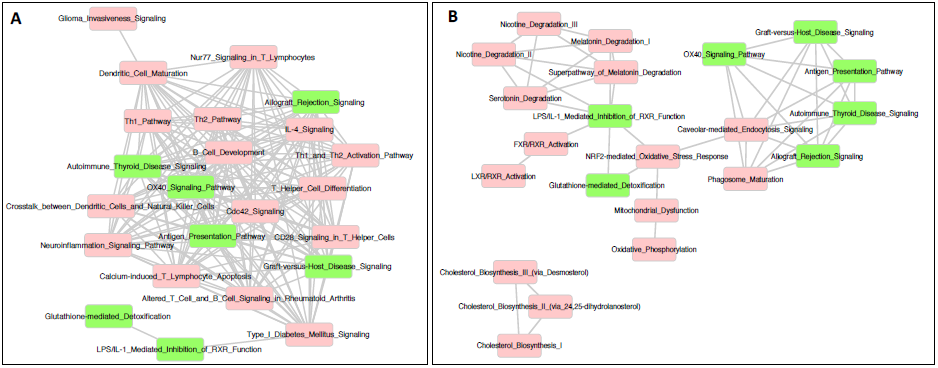
Ingenuity pathway analysis of significant and interconnected eQTLs in (**A**) glomerulus and (**B**) tubulointerstitium. Pathways shared in GLOM and TI are in green. Distinct pathways are in red.

Among the most significant GLOM-specific eQTLs as computed by eQtlBma, were those of notable relevance to NS, particularly with phospholipase C gamma 2 (*PLCG2*) and vacuolar protein sorting 33b (*VPS33B*). *PLCG2* has been implicated in pediatric SSNS via genome-wide rare-variant association study^7^. *VPS33B* is a vacuolar protein and functions in vesicle mediated protein sorting, predominantly in the late endosome/lysosome^44^. Importantly, mutations in *VPS33B* cause **A**rthrogryposis, **r**enal dysfunction, and **c**holestasis (ARC) Syndrome,” in which the renal phenotype includes NS ^45-47^.

### Cell-type specificity of significant GLOM & TI eQTLs

There is substantial cellular heterogeneity within the GLOM and TI which could potentially confound discovery of eQTLs that are cell specific or restricted to a subset of cells. We hypothesized that we could deconvolute the GLOM and TI eQTLs into specific cell-types through use of independent single-cell RNA-Seq data of human kidney. Because scRNA-Seq on NEPTUNE research biopsy cores is not feasible due to a lack of sufficient starting material, we integrated our eQTL data with single-cell RNA-Seq data derived from healthy portions of adult tumor nephrectomy tissue.

Single cell transcriptome analyses of 4,734 cells identified 14 clusters of specific cell types as defined by differentially expressed cell-type specific genes (**Figure 5, Supplemental Table 3)**. The number of cells per cluster ranged from 49 in the podocyte to 1712 within the proximal tubule. For example, there were 772 and 973 genes that were significantly differentially expressed in the podocyte and proximal tubule cell clusters, respectively.

**Table 3:** Top GLOM and TI genes associated with IgA Nephropathy case status (FDR < 0.1) via transcriptome-wide association study

| Tissue | Gene | FDR | Direction of expression associated w/IgAN cases |
| --- | --- | --- | --- |
| Glomerulus | *HLA-DRB5* | 8.29E-06 | ↓ |
|  | *RNF5* | 2.24E-05 | ↓ |
|  | *HLA-DPB1* | 2.24E-05 | ↑ |
|  | *HLA-DQA1* | 3.57E-05 | ↑ |
|  | *HLA-DPA1* | 4.82E-04 | ↑ |
|  | *HLA-DQA2* | 6.95E-04 | ↑ |
|  | *HLA-DQB1* | 1.12E-02 | ↓ |
|  | *HLA-DQB2* | 4.11E-02 | ↓ |
|  | *BTN3A2* | 5.06E-02 | ↑ |
|  | *ZKSCAN8* | 5.06E-02 | ↓ |
|  | *HLA-DRB1* | 5.10E-02 | ↓ |
|  | *CRYBA2* | 7.72E-02 | ↑ |
|  | *C2orf74* | 7.72E-02 | ↓ |
| Tubulointerstitium | *HLA-DRB5* | 3.29E-09 | ↓ |
|  | *HLA-DQB2* | 2.57E-05 | ↓ |
|  | *HLA-DQB1* | 1.21E-04 | ↓ |
|  | *HLA-DRB1* | 6.60E-04 | ↓ |
|  | *HLA-DPB1* | 1.19E-03 | ↑ |
|  | *ATF6B* | 3.01E-03 | ↑ |
|  | *HLA-DPA1* | 3.59E-03 | ↑ |
|  | *VWA7* | 1.85E-02 | ↓ |
|  | *SERPINA4* | 8.31E-02 | ↓ |
|  | *LAMB4* | 8.77E-02 | ↑ |
|  | *HIST1H4L* | 9.17E-02 | ↑ |
|  | *C2orf74* | 9.17E-02 | ↓ |

**Figure 5:**
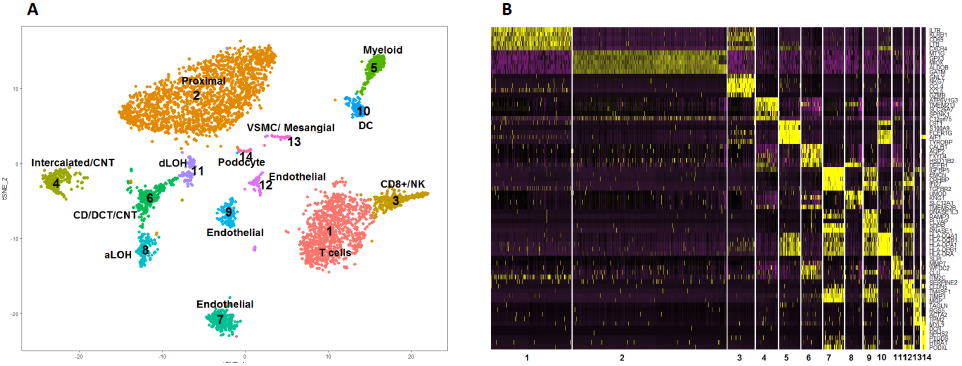
Results of adult kidney single cell RNA-Seq experiment of 4,734 cells resolved into 14 clusters. (**A**) tSNE plot of 14 kidney cell types; (**B**) heat map show the top 5 cell-type specific gene expression for each of the 14 clusters

As shown in **Figure 6**, GLOM and TI eQTL genes were most significantly enriched in podocyte (OR: 2.5, p=8 ×10^−11^) and proximal tubule cells (OR: 3.4, p=1 ×10^−43^), respectively. We note that the GLOM eQTLs are second most enriched in proximal cells. We believe that this is a function of the microdissection process, in which we are unable to remove all proximal tubular cells from the glomeruli. Assignations of GLOM and TI eQTLs to specific cell-type clusters is provided in **Supplementary Table 4.**

**Figure 6:**
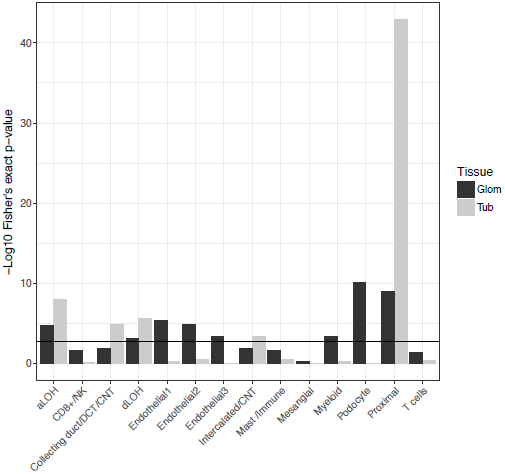
Enrichment of GLOM and TI eQTLs in specific kidney cell types. A Fisher’s exact test was used to determine whether transcripts significantly enriched in one of 14 specific cell types from single cell RNA-Seq studies of adult kidney were enriched for GLOM and/or TI eQTLs. The horizontal line represents the Bonferroni corrected threshold for significance. We observe that GLOM eQTLs were most enriched in podocytes while TI eQTLs were most enriched in proximal tubular cell types.

We used Genomatix to annotate the 79 podocyte-specific GLOM eQTLs. The top three biologic processes (all p< 7 ×10^−07^) were “vesicle-mediated transport”, “endocytosis”, and “regulation of locomotion”, the top molecular function was “extracellular matrix binding” (P<7 ×10^−06^), and the top signal transduction pathway was “integrin signaling” (p< 5 ×10^−05^).

### Kidney tissue-based TWAS for IgAN

Using an strategy based on the prediXcan^42^ approach with minor modifications to accommodate microarray data, we predicted gene expression with a cross-validated r^2^ > 0.01 (correlation > 0.1) in 3294 genes from the glom and 3869 genes from the tub. 39% of glom models and 42% of tub models had a cross-validated correlation > 0.2 **(Supplementary Table 5)**. Among prediction models with r^2^ > 0.01, 73% of models in the glom and tub had at least 10 variants selected in the prediction models. The number of variants selected in the model does not correspond to the number of independent eQTLs because the elastic net has a grouping effect where correlated variables tend to selected or not as a group^48^. This results in linked variants being selected in the prediction model with the effect size distributed across linked variants.

We discovered 13 GLOM and 12 TI eQTLs associated with IgAN with a FDR < 0.1. In both tissues, decreased expression of *HLA-DRB5* was the most strongly associated with IgAN (FDR=8.3 x 10^−6^ in GLOM & 3.3 x 10^−9^ in TI) (**Table 3 & Supplemental Figure 3**).

### Characterizing known kidney GWAS alleles

We hypothesized that GLOM and TI eQTLs could be used to enhance interpretations of known GWAS alleles beyond those for nephrotic syndrome or other glomerular diseases. For example, in one of the first GWAS of chronic kidney disease (CKD), the lead SNP was rs12917707, 3.6 kb upstream of uromodulin (*UMOD*)^49^, a finding that has been replicated numerous times and has led to an entire field of study regarding *UMOD* in CKD. rs12917707 is not an eQTL across 44 tissues in GTEx (http://gtexportal.org). By contrast, it is a significant TI eQTL in our data, with the risk allele associated with increased expression of *UMOD* (**Supplementary Figure 4**). Our TI results are concordant with reports that the risk allele is associated with increased UMOD transcript expression in tumor nephrectomy tissue^50^ and increased urinary protein UMOD expression^51^.

## Discussion

As shown elsewhere, biologic insights from eQTL studies becomes particularly powerful when using the tissues or cells directly involved in the disease, such as adipose and muscle eQTLs for Type 2 Diabetes (T2D)^52^, leukocytes in diverse immune-mediated diseases^53^ ^54^, adipocytes in obesity^55^, or brain in schizophrenia^56^. Here, we used a *cis*-eQTL study design to discover genetic variants associated with steady-state mRNA levels of genes expressed in the glomeruli and tubulointerstitium of nephrotic syndrome patients. Through the use of eQtlBma and the integration of single-cell RNA-Seq data, we gained greater specificity with regards to compartment and/or cell specificity of the eQTLs we discovered.

At a genome-wide level of significance, we discovered 894 GLOM eQTLs and 1767 TI eQTLs in 136 and 166 eligible patients, respectively. By using DAP, we could fine-map these eQTLs, identifying specific SNVs with the highest probability of driving the signal, and also discovering a subset of GLOM and TI eQTL whose association is most likely to be driven by multiple, independent loci. Moving forward, narrowing the sets of implicated SNVs to identify the one that is causal, and understanding the genome biology underlying its impact on regulation will be important as we seek to discover targets and mechanism upon which we can intervene therapeutically. Given the tissue- and cell-specific nature of genomic regulatory elements, the publication and sharing of epigenetic data derived from kidney and GLOM and TI cells will be vital.

When functionally annotating the DAP-significant GLOM and TI eQTLs, we observed an enrichment of immunity functions and pathways in both tissues, a not unexpected result given the known role of these pathways in NS. The cellular origin of these eQTLs are unclear, as podocytes can express these genes^57^, there are resident immune cells in the kidney^58^, and a presence of circulating immune cells. Future studies that compare eQTLs in circulating immune cells to those in the kidneys from the same patient may inform us of the contribution of each in patients with NS. If we can ultimately link these eQTLs to clinical phenotypes, this may provide an opportunity to target the mechanism or the gene’s expression level for therapeutics.

eQtlBma and single-cell RNA-Seq data provided additional insights and specificity to these studies. The GLOM- and TI-specific eQTLs identified by eQtlBma were enriched for processes related to the normal physiology of these compartments rather than the immune-related functions shared across tissues. Using the scRNA-Seq data, we found that GLOM eQTLs were most enriched in podocytes, TI eQTLs were most enriched in the proximal tubule, and the majority of eQTLs could be assigned to a cell-type. These findings should provide confidence in broadening the use of these data beyond NS discovery.

Through a TWAS of IgAN and a single SNP lookup of the lead SNP at the CKD-associated UMOD locus, we demonstrated the utility of these eQTL data to gain further insights from existing GWAS of kidney disease. We focused on IgAN because, among glomerular disorders, it has the largest GWAS readily available. In this way, we identified a limited number of genes in which change in genetically predicted expression was associated with the disease status. Notably, decreased expression of *HLA-DRB5* was the most significantly associated with IgAN in our TWAS in both in both GLOM and TI compartments. This suggests potential protective role of DRB5 gene in IgAN, although we recognize that the extended region of linkage disequilibrium across the HLA region limits the interpretability of this finding. We also appreciate that the causal cells for IgAN may also reside outside the kidney, with multiple hits across cell types needed for disease pathogenesis. Future studies that perform eQTLs across these cell types should be further illuminating. As larger GWAS datasets for other primary glomerular disorders, such as FSGS, MN, or MCD, become available, the TWAS approach should become even more relevant.

The inclusion criteria for patients enrolled in NEPTUNE was a need for a kidney biopsy for suspicion of primary NS due and there was no limitation to specific histologic diagnoses, eGFR, or response to immunosuppression. As such, this study should be interpreted in this light; namely that we are identifying eQTLs that are observed in the kidneys of patients with proteinuric glomerular filtration barrier failure. However, we found computational and experimental strategies can be applied to GLOM and TI eQTLs data to derive insights that are broadly relevant to underlying biology of these structures and specific to particular cell-types. Future GLOM and TI eQTL studies in normal kidneys and those with a specific histologic or molecular diagnosis (and of similar sample sizes) would complement this study with insights about similarities and differences in the genetics of gene expression as a function of disease state.

In our opinion, sharing these eQTL data in an easily accessible manner is as equally important as any of the specific discoveries that we report here. To this end, we have added a stand-alone eQTL browser “nephQTL”, http://nephqtl.org, to our existing NephVS software (http://nephvs.org). nephQTL has a searchable browser of the summary-level MatrixEQTL and DAP output for GLOM and TI, with both summary statistics and visualizations of the eQTLs. The full MatrixEQTL output is also available for download and secondary use. Our hope is that unrestricted access to this unique database will be useful to the wider community, catalyzing and accelerating discoveries that will ultimately lead to improved health for patients with NS and beyond.

### Description of Supplemental Data

The Supplemental Data consists of 5 figures and 5 tables/files.

## Web Resources

Nephrotic syndrome variant server (http://nephvs.org)

NephQTL browser (http://nephQTL.org)

## Conflicts of Interest

There are no conflicts of interest to declare.

## Acknowledgments

M.G.S. is supported by the Charles Woodson Clinical Research Fund, the Ravitz Foundation, and by National Institutes of Health RO1-DK108805

KK is supported by National Institutes of Health RO1-DK105124

The Nephrotic Syndrome Study Network Consortium (NEPTUNE), U54-DK-083912, is a part of the National Center for Advancing Translational Sciences (NCATS) Rare Disease Clinical Research Network (RDCRN), supported through a collaboration be-tween the Office of Rare Diseases Research (ORDR), NCATS, and the National Institute of Diabetes, Digestive, and Kidney Diseases. RDCRN is an initiative of ORDR, NCATS. Additional funding and/or programmatic support for this project has also been provided by the University of Michigan, NephCure Kidney International, and the Halpin Foundation.

## Members of the Nephrotic Syndrome Study Network (NEPTUNE)

*NEPTUNE Enrolling Centers*

*Case Western Reserve University, Cleveland, OH*: J Sedor^*^, K Dell**, M Schachere^#^

*Children’s Hospital, Los Angeles, CA*: K Lemley^*^, L Whitted^#^

*Children’s Mercy Hospital, Kansas City, MO*: T Srivastava^*^, C Haney^#^

*Cohen Children’s Hospital, New Hyde Park, NY:* C Sethna^*^, K Grammatikopoulos^#^

*Columbia University, New York, NY:* G Appel^*^, M Toledo^#^

*Emory University, Atlanta, GA:* L Greenbaum^*^, C Wang**, B Lee^#^

*Harbor-University of California Los Angeles Medical Center:* S Adler^*^, C Nast^*‡^, J La Page^#^

*John H. Stroger Jr. Hospital of Cook County, Chicago, IL:* A Athavale^*^

*Johns Hopkins Medicine, Baltimore, MD:* A Neu^*^, S Boynton^#^

*Mayo Clinic, Rochester, MN:* F Fervenza^*^, M Hogan**, J Lieske^*^, V Chernitskiy^#^

*Montefiore Medical Center, Bronx, NY:* F Kaskel*, N Kumar*, P Flynn#

*NIDDK Intramural, Bethesda MD:* J Kopp^*^, E Castro-Rubio^#^, J Blake^#^

*New York University Medical Center, New York, NY:* H Trachtman^*^, O Zhdanova**, F Modersitzki^#^, S Vento^#^

*Stanford University, Stanford, CA:* R Lafayette^*^, K Mehta^#^

*Temple University, Philadelphia, PA:* C Gadegbeku^*^, D Johnstone**

*University Health Network Toronto:* D Cattran^*^, M Hladunewich**, H Reich**, P Ling^#^, M Romano^#^

*University of Miami, Miami, FL:* A Fornoni^*^, L Barisoni^*^, C Bidot^#^

*University of Michigan, Ann Arbor, MI:* M Kretzler^*^, D Gipson*, A Williams^#^, R Pitter^#^

*University of North Carolina, Chapel Hill, NC:* P Nachman^*^, K Gibson*, S Grubbs^#^, A Froment^#^

*University of Pennsylvania, Philadelphia, PA:* L Holzman^*^, K Meyers**, K Kallem^#^, FJ Cerecino^#^

*University of Texas Southwestern, Dallas, TX:* K Sambandam^*^, E Brown**, N Johnson^#^

*University of Washington, Seattle, WA:* A Jefferson^*^, S Hingorani**, K Tuttle**^§,^ L Curtin^#^, S Dismuke^#^, A Cooper^#§^

*Wake Forest University, Winston-Salem, NC:* B Freedman^*^, JJ Lin**, M Spainhour^#^, S Gray^#^

*Data Analysis and Coordinating Center*: M Kretzler, L Barisoni, C Gadegbeku, B Gillespie, D Gipson, L Holzman, L Mariani, M Sampson, P Song, J Troost, J Zee, E Herreshoff, C Kincaid, C Lienczewski, T Mainieri, A Williams

*National Institute of Diabetes and Digestive and Kidney Diseases (NIDDK) Program Office:* K Abbott, C Roy

*The National Center for Advancing Translational Sciences (NCATS) Program Office:* T Urv, PJ Brooks

*Principal Investigator; **Co-investigator; ^#^Study Coordinator

^‡^Cedars-Sinai Medical Center, Los Angeles, CA

^§^Providence Medical Research Center, Spokane, WA

## Supplementary Figures

**Supplementary Figure 1**: Genotype-based Principal Components 1 & 2 of the 187 participants in the eQTL study.

**Supplementary Figure 2:** Expected model size (# of independent SNVs) for eQTLs in GLOM and TUB, as predicted by DAP.

**Supplementary Figure 3:** A significant StringDB-derived, protein-protein interaction network emerging from analysis of the GLOM-specific eQTLs as computed by eQtlBma

**Supplementary Figure 4:** QQ plot of the results from the metaXscan analysis of summary statistics from the IgA nephropathy GWAS with our glomerular eQTL data

**Supplementary Figure 5:** Tubulointerstitial matrix eQTL output for rs12917707, the lead GWAS SNP for CKD published by the CKDGen consortium. The risk allele “G” is associated with increased *UMOD* transcript expression, which replicates independent studies in bulk renal cortex and urinary UMOD protein expression.

## Supplemental Tables

**Supplementary Table 1:** List of significant eQTL from GLOM and TI discovered using DAP. Significance defined as an FDR <0.05.

**Supplementary Table 2:** GLOM and TI specific eQTLs via eQtlBma

**Supplementary Table 3:** 99 GLOM eQTLs that are podocyte-specific

**Supplementary Table 4:** GLOM and TI eQTLs whose genes’ expression could be confidently predicted using MetaXscan

**Supplementary Table 5**: Results of TWAS of IgA Nephropathy using GLOM and TI-eQTL.

**Figure S1:**
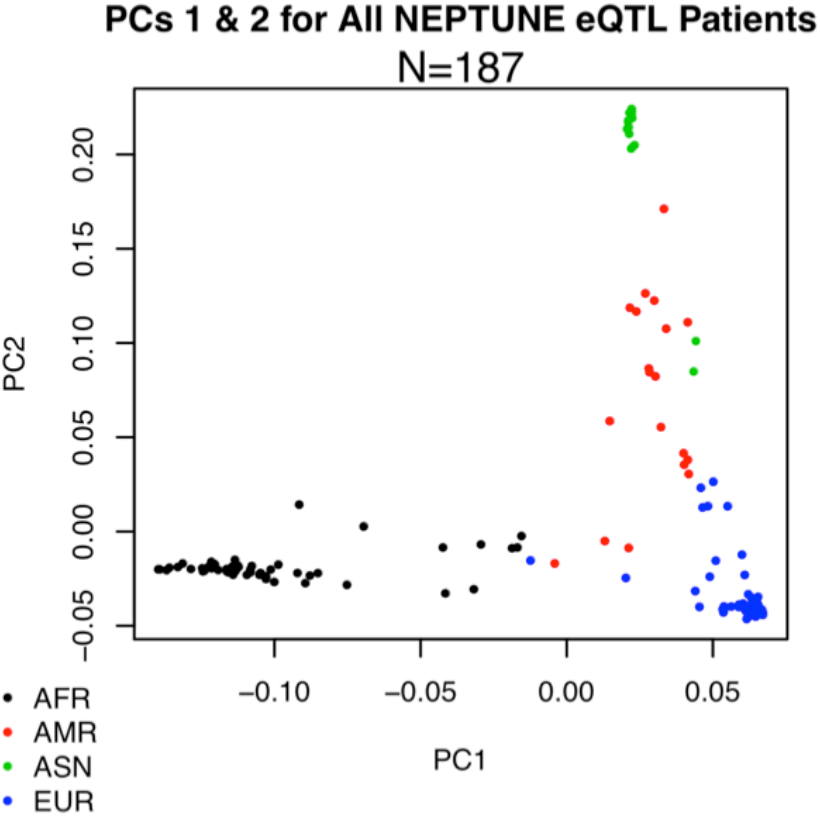
Genotype-based Principal Components 1 & 2 of the 187 participants in the eQTL study.

**Figure S2:**
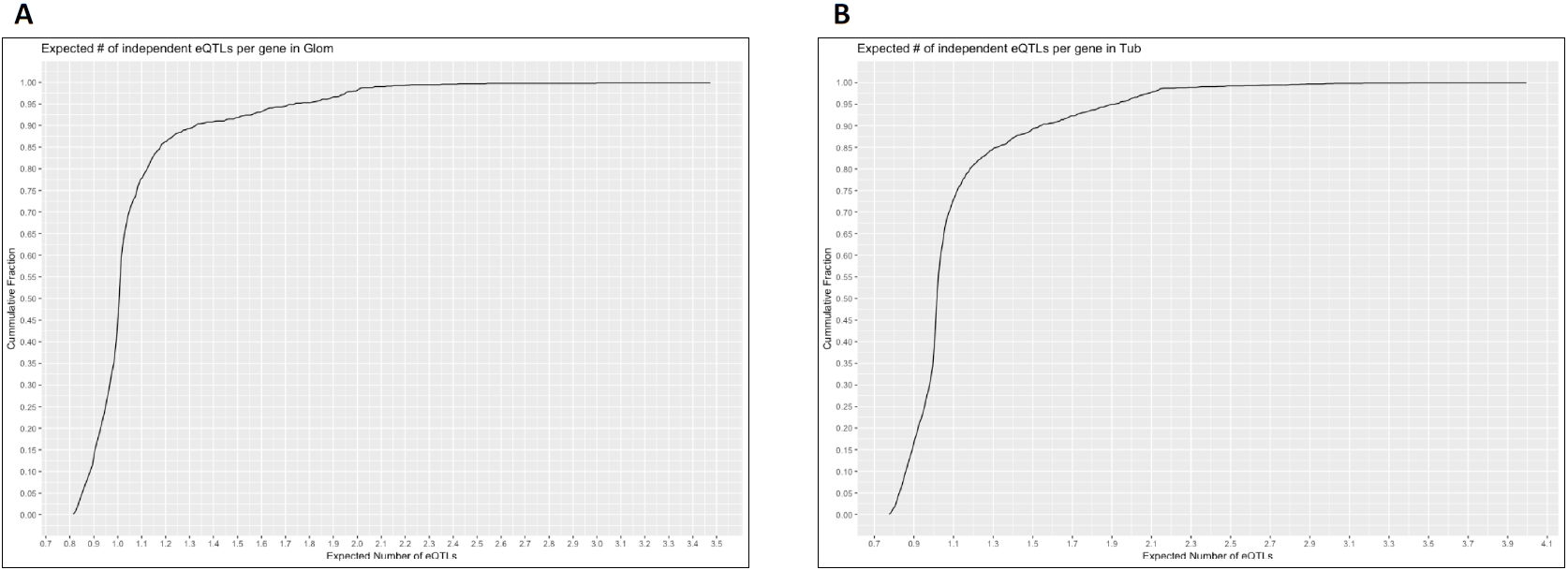
Expected model size (# of independent SNVs) for eQTLs in GLOM and TUB, as predicted by DAP.

**Figure S3:**
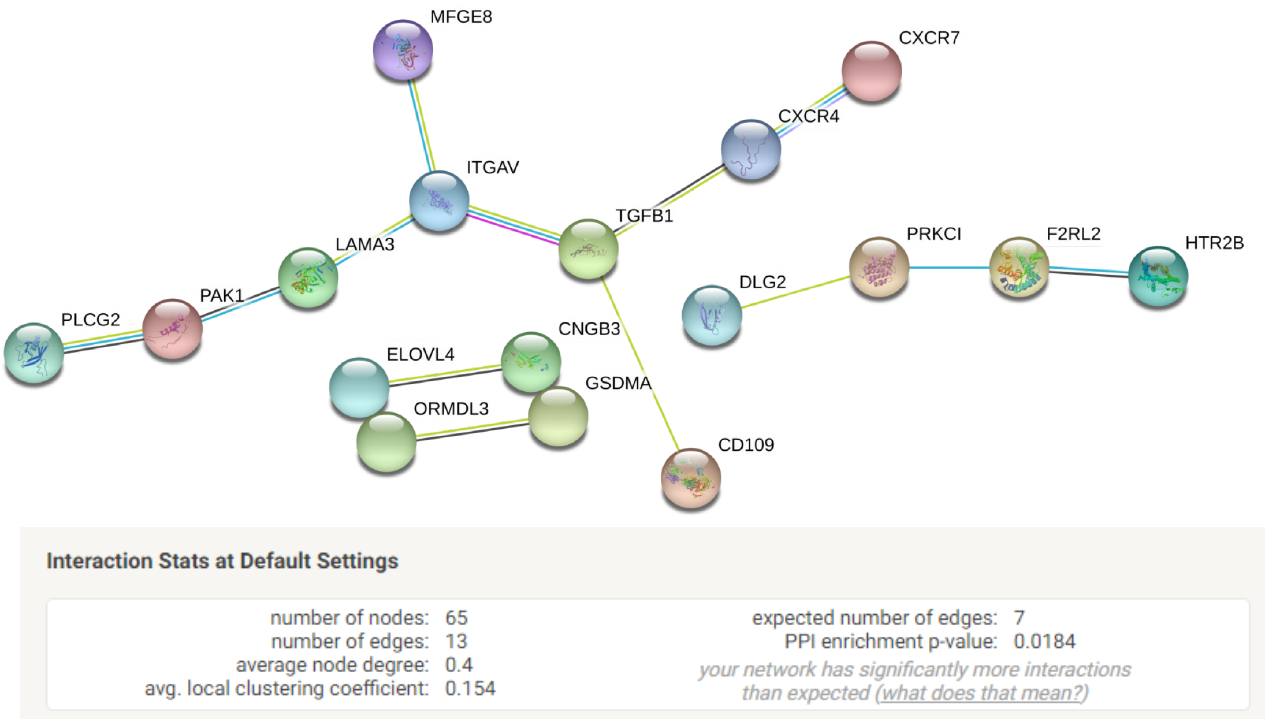
A significant StringDB-derived, protein-protein interaction network emerging from analysis of the GLOM-specific eQTLs as computed by eQtlBma

**Figure S4:**
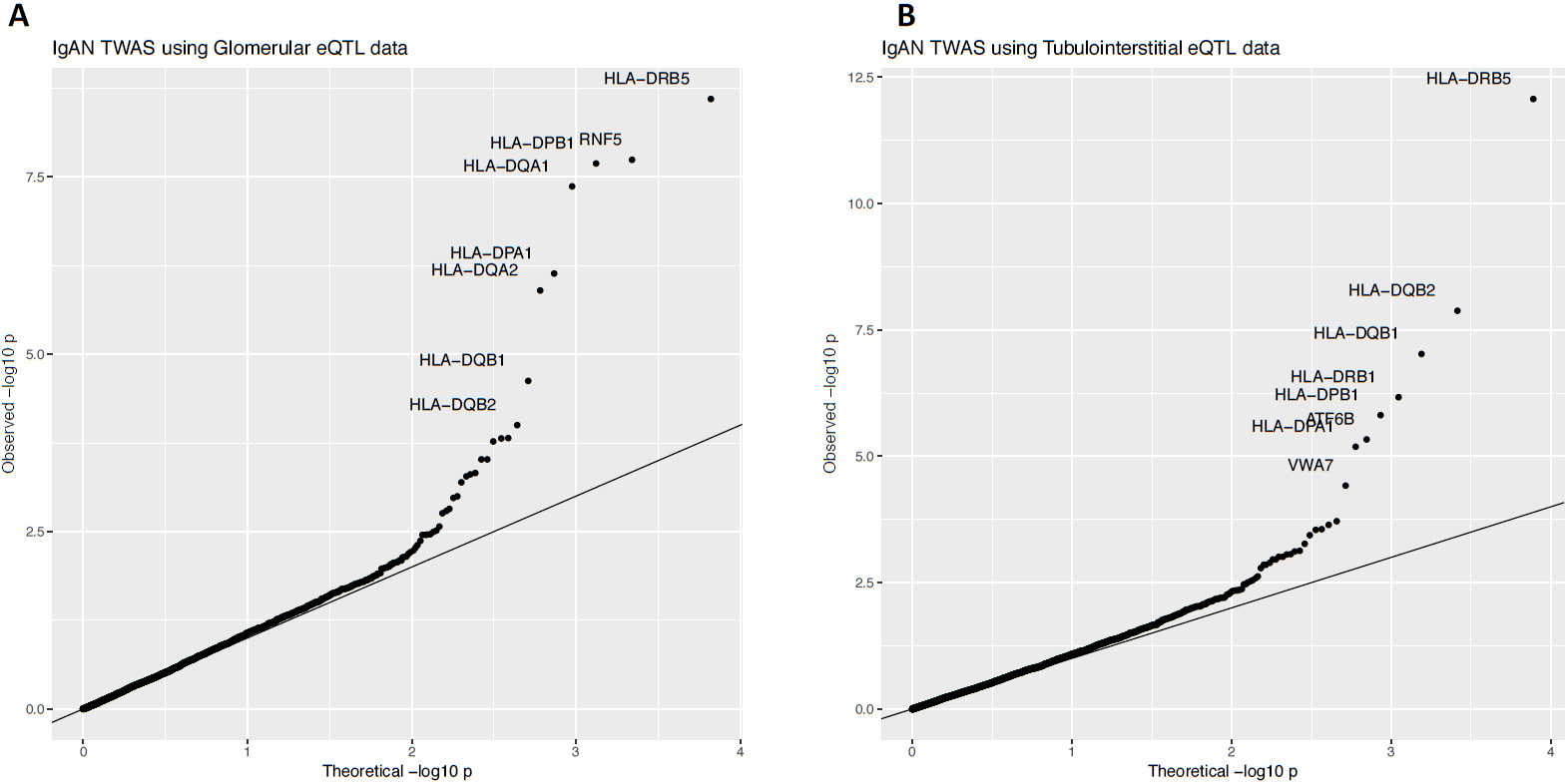
QQ plot of the results from the metaXscan analysis of summary statistics from the IgA nephropathy GWAS with our (**A**) glomerular and (**B**) tubulointerstitial eQTL data

**Figure S5:**
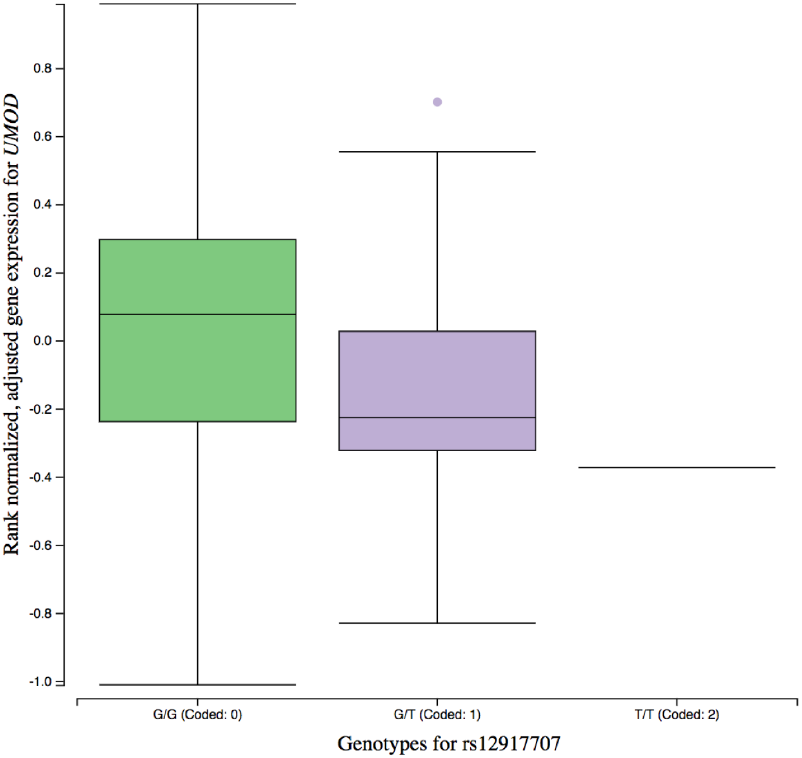
Tubulointerstitial matrix eQTL output for rs12917707, the lead GWAS SNP for CKD published by the CKDGen consortium. The risk allele “G” is associated with increased *UMOD* transcript expression, which replicates independent studies in bulk renal cortex and urinary UMOD protein expression.

